# Atovaquone-induced therapeutic rewiring of melanoma metabolism

**DOI:** 10.1101/2025.09.08.674005

**Authors:** Mayra Betancourt Ponce, Alexa R. Heaton, Samantha K. Burkard, Aurora D’Amato, Nicha Boonpattrawong, Melissa C. Skala, Manish Patankar, Lisa Barroilhet

**Affiliations:** University of Wisconsin School of Medicine and Public Health, Department of Obstetrics and Gynecology, Madison, WI; Morgridge Institute for Research, Madison, WI; University of Wisconsin, Department of Biomedical Engineering, Madison WI; University of Wisconsin, Department of Human Oncology, Madison WI; University of Wisconsin, Department of Computer Sciences, Madison WI; University of Wisconsin, Global Health Institute, Madison WI

## Abstract

Melanoma continues to be the deadliest form of skin cancer, emphasizing the need for new therapeutic strategies. Targeting tumor metabolism, particularly oxidative phosphorylation (OXPHOS) has emerged as a promising approach due its role in melanoma tumor survival, metastasis, and treatment resistance. In this study, we investigate the metabolic and antitumor effects of atovaquone, an FDA-approved and safe OXPHOS inhibitor that has not been previously tested in melanoma. We show that atovaquone rapidly and effectively inhibits OXPHOS and impairs glycolysis in melanoma, leading to metabolic reprogramming observed via metabolic imaging and marked depletion in energy (ATP) stores. Atovaquone also induces oxidative stress, evidenced by increased reactive oxygen species levels, DNA damage, and upregulation of antioxidant proteins. Moreover, atovaquone reduced melanoma cell viability and migration *in vitro*, and slowed tumor growth *in vivo*. Notably, these effects were observed in both *BRAF*-wild-type and mutant melanoma models, suggesting its potential as an effective treatment across different subtypes. Our study identifies atovaquone as a metabolic disruptor with antitumor activity against melanoma, supporting further investigation as a repurposed therapeutic strategy.

## INTRODUCTION

Melanoma is the deadliest form of skin cancer.^1,2^ Even though it only accounts for 4% of skin cancer cases, it is responsible for 75% of skin cancer-related deaths.^3^ The current standard of care consists of surgical excision, which is most effective when disease is diagnosed in early, localized stages.^1,3^ In advanced, disseminated stages, surgery is typically combined with immunotherapy or targeted therapy depending on patients’ mutational profiles.^1,3^ While these adjuvant therapies have improved outcomes in subsets of patients, many patients are non-responsive and develop resistance.^4–6^ With the rising incidence of melanoma,^2^ there is a critical need for improved treatment strategies.

Oxidative phosphorylation (OXPHOS) has emerged as an important contributor to melanoma bioenergetics and a promising therapeutic target. Although proliferating melanoma tumors are often characterized by glycolytic metabolic phenotypes, OXPHOS has also been shown to play key roles in tumor survival, metastasis, and treatment resistance.^7–10^ Several OXPHOS inhibitors have shown potential preclinically but have failed to succeed in clinical trials. For example, the electron transport chain (ETC) complex I inhibitors BAY87-2243 and IACS-010759 have been reported to have activity against melanoma.^11,12^ Similarly, metformin, which also targets ETC complex I, reduces melanoma cell viability and enhances sensitivity to other treatments such as platinum drugs and checkpoint inhibitors.^13–15^ While the clinical trials evaluating these complex I inhibitors as cancer therapies have not specifically focused on melanoma, they have highlighted challenges, including high toxicity with BAY87-2243 and IACS-010759, or lack of effectiveness with metformin.

In this study, we investigate for the first time the effects of atovaquone, an alternative OXPHOS inhibitor, in melanoma. Atovaquone is a well-tolerated, orally available, FDA-approved, ETC complex III inhibitor that has shown efficacy in different cancer types.^16–19^ Given that melanoma cells often retain functional OXPHOS machinery that can contribute to tumor progression,^7,20^ we hypothesized that atovaquone would disrupt melanoma metabolism and exhibit antitumor activity. Our findings demonstrate that atovaquone effectively inhibits OXPHOS, induces oxidative stress, and suppresses *in vitro* and *in vivo* melanoma cell growth across different melanoma models. These results establish a strong foundation for further evaluation of atovaquone as a therapeutic agent for melanoma.

## RESULTS

### Atovaquone inhibits OXPHOS in melanoma

Given that atovaquone has not been previously studied in the context of melanoma, we first verified its metabolic effects on melanoma cells using Seahorse assays. Atovaquone significantly reduced oxygen consumption rates (OCR) to baseline levels as early as 1-hour with effects sustained at 24 hours post-treatment in B16 and B78 cells (**Figure 1A-B**). These two well-established cell lines have distinct intrinsic characteristics; B78 is an amelanotic line derived from the melanin-producing B16 cells.^21^

**Figure 1.**
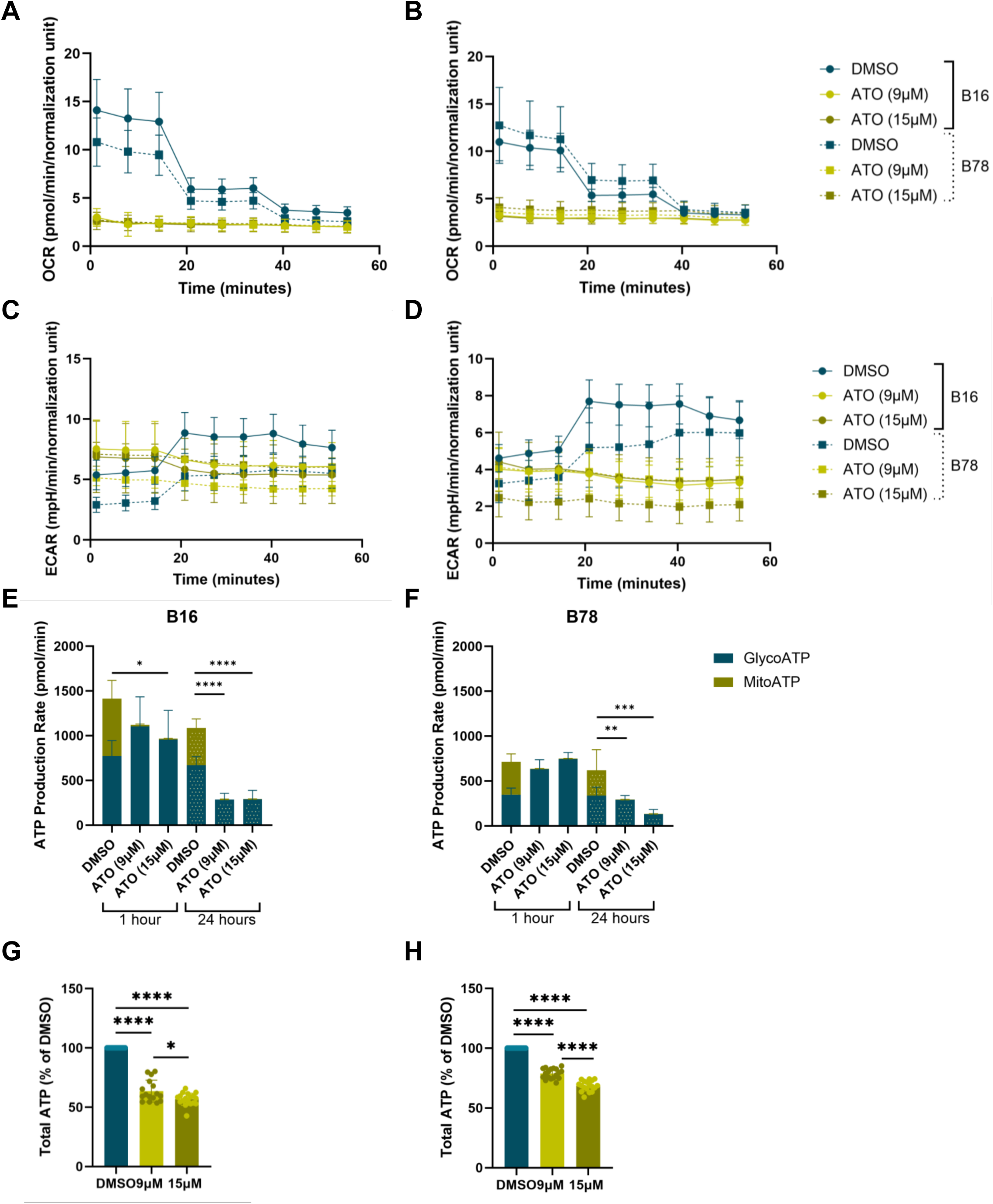
Atovaquone disrupts melanoma cell metabolism and ATP production. Seahorse XF Real-Time ATP Rate Assay was performed at 1 and 24 hours post-atovaquone treatment. Oxygen consumption rates were significantly decreased in atovaquone-treated cells compared to vehicle (DMSO)-treated cells at 1 (A) and 24 hours post-treatment (B). Extracellular acidification rates were increased in atovaquone-treated cells compared to vehicle (DMSO)-treated cells at 1 hour (C), while unchanged or decreased at 24 hours post-treatment (D). Total ATP production rates were decreased in atovaquone-treated cells compared to vehicle at 24 hours post-treatment in both B16 (E) and (B78 cells (F). Total ATP levels were significantly decreased in atovaquone-treated cells compared to vehicle (DMSO)-treated cells at 24 hours in both B16 (E) and (B78 cells (F), as observed via CellTiter-Glo® 2.0 assay. Similar results were observed in B16 and B78 cell lines. \**P*<0.05, \*\**P*<0.01, \*\*\**P*<0.001, \*\*\*\**P*<0.0001.

We next calculated the contribution of glycolysis and OXPHOS to ATP production (**Supplemental Table 1**). In untreated B16 cells, 55.1% and 62.1% of ATP production was attributed to glycolysis (GlycoATP) whereas 44.9% and 37.9% was attributed to OXPHOS (mitoATP) at 1 and 24 h timepoints, respectively. In contrast, atovaquone-treated cells exhibited a shift where >98.5% was attributed to glycolysis while <1.5% was attributed to OXPHOS, regardless of the treatment concentration (9 and 15 μM) or time point (1 and 24 hours). Consequently, ATP rate indexes, defined as the ratio of mitochondrial to glycolytic ATP production, were significantly decreased from 0.82 in vehicle-treated cells to less than 0.016 in atovaquone-treated cells at 1 hour and from 0.61 to <0.010 in atovaquone-treated cells at 24 hours. Similar results were observed in B78 cells. These results confirm that atovaquone robustly inhibits OXPHOS in B16 and B78 melanoma cells.

Given these effects, we anticipated a compensatory increase in extracellular acidification rates (ECAR), which assess glycolytic activity. Interestingly, we observed an initial increase in ECAR at the 1-hour time point, but these effects were abrogated at 24 hours, where ECAR levels were comparable to or even lower than vehicle controls (**Figure 1C-D**).

This inability to sustain glycolytic compensation, along with OXPHOS inhibition, was reflected in ATP production rates, which were significantly reduced in atovaquone-treated cells compared to vehicle, especially at 24 hours when both OCR and ECAR were decreased (**Figure 1E-F**). ATP-based viability assays further showed significantly decreased total ATP levels in atovaquone-treated compared to vehicle at 24 hours (**Figure 1G-H**).

To further assess metabolic fluctuations following atovaquone treatment, we performed multiphoton optical metabolic imaging in B78 cells. Multiphoton imaging allows us to visualize the endogenous metabolic coenzymes NAD(P)H and FAD. Utilizing their unique autofluorescence spectra and fluorescence lifetimes, we measured optical redox ratios (ORR), which provide an indirect assessment of the redox balance of the cell and protein-binding changes with atovaquone treatment.^21,22^ At 1 hour, ATO-treated cells exhibited a significant increase in free NAD(P)H (α_1_), consistent with an initial increase in glycolysis (**Figure 2C**). At 24 hours, ATO-treated cells exhibited significantly decreased NAD(P)H intensity and decreased free NAD(P)H (α_1_) compared to vehicle, consistent with a decrease in glycolysis (**Figure 2F,G**). These metabolic changes mirror the Seahorse data (**Figure 1**) that shows an initial glycolytic ECAR increase at 1 hour, followed by a sharp decrease in ECAR at 24. These changes may result from variation in NAD(P)H production through pathways like glycolysis, or increased NAD(P)H consumption via antioxidant mechanisms. At 24 hours, we also observed a significant increase in free FAD (α_2_) following atovaquone treatment (**Figure 2H**), possibly due to ETC complex III inhibition by atovaquone and a resulting buildup of oxidized FAD. Ultimately, we observed a significant decrease in the ORR of atovaquone-treated cells compared to vehicle controls at 24 hours (**Figure 2I**), suggesting a shift toward an oxidized state and reflecting the decrease in NAD(P)H intensity (**Figure 2F**). This may also be consistent with oxidative stress. The 24-hour atovaquone induced changes trend the same as other ETC complex III inhibitors such as antimycin A.^23^ As with our Seahorse data, these results show that atovaquone disrupts melanoma cell metabolism beyond OXPHOS inhibition.

**Figure 2.**
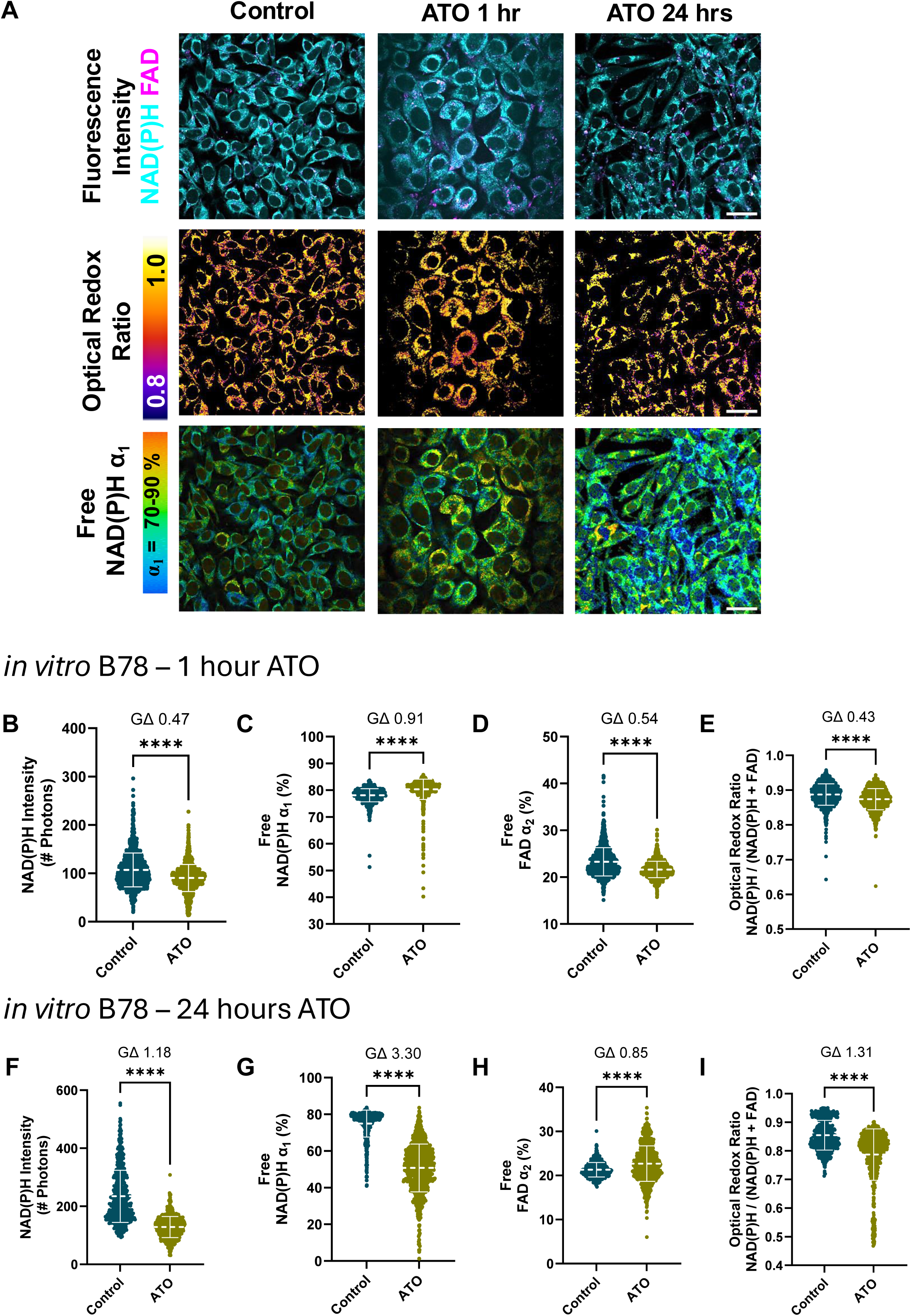
Optical metabolic imaging shows *in vitro* B78 metabolic changes during atovaquone treatment. (A) Representative *in vitro* B78 images showed fluorescence intensity changes of metabolic coenzymes NAD(P)H and FAD (top), optical redox ratio changes (middle), and NAD(P)H fluorescence lifetime changes (bottom) over time with atovaquone treatment (scale bar 50 μm). (B-E) Quantified single cell B78 metabolic changes with 1 hour atovaquone treatment (15 μM, *n* = 1276 pre-treatment control cells, *n* = 1137 post-treatment atovaquone treated cells). (F-I) Quantified single cell B78 metabolic changes with 24 hour atovaquone treatment (15 μM, *n* = 573 pre-treatment control cells, *n* = 827 post-treatment atovaquone cells). \*\*\*\**P* < 0.0001, Glass’s Δ effect size, 2 biological replicates each, 2 technical replicates each, mean ± SD.

### Atovaquone induces oxidative stress in melanoma cells

OXPHOS disruption leads to the accumulation of reactive oxygen species (ROS).^24^ To assess whether atovaquone increases ROS in melanoma, we quantified intracellular ROS following atovaquone treatment using the cell-permeable compound H_2_DCFDA. Once inside the cell, H_2_DCFDA is deacetylated into a non-fluorescent form that can be oxidized by ROS to generate a fluorescent, quantifiable signal.^25^ Following atovaquone treatment, ROS levels were elevated as early as 30 minutes and increased significantly by 24 hours compared to controls (**Figure 3A-B**).

**Figure 3.**
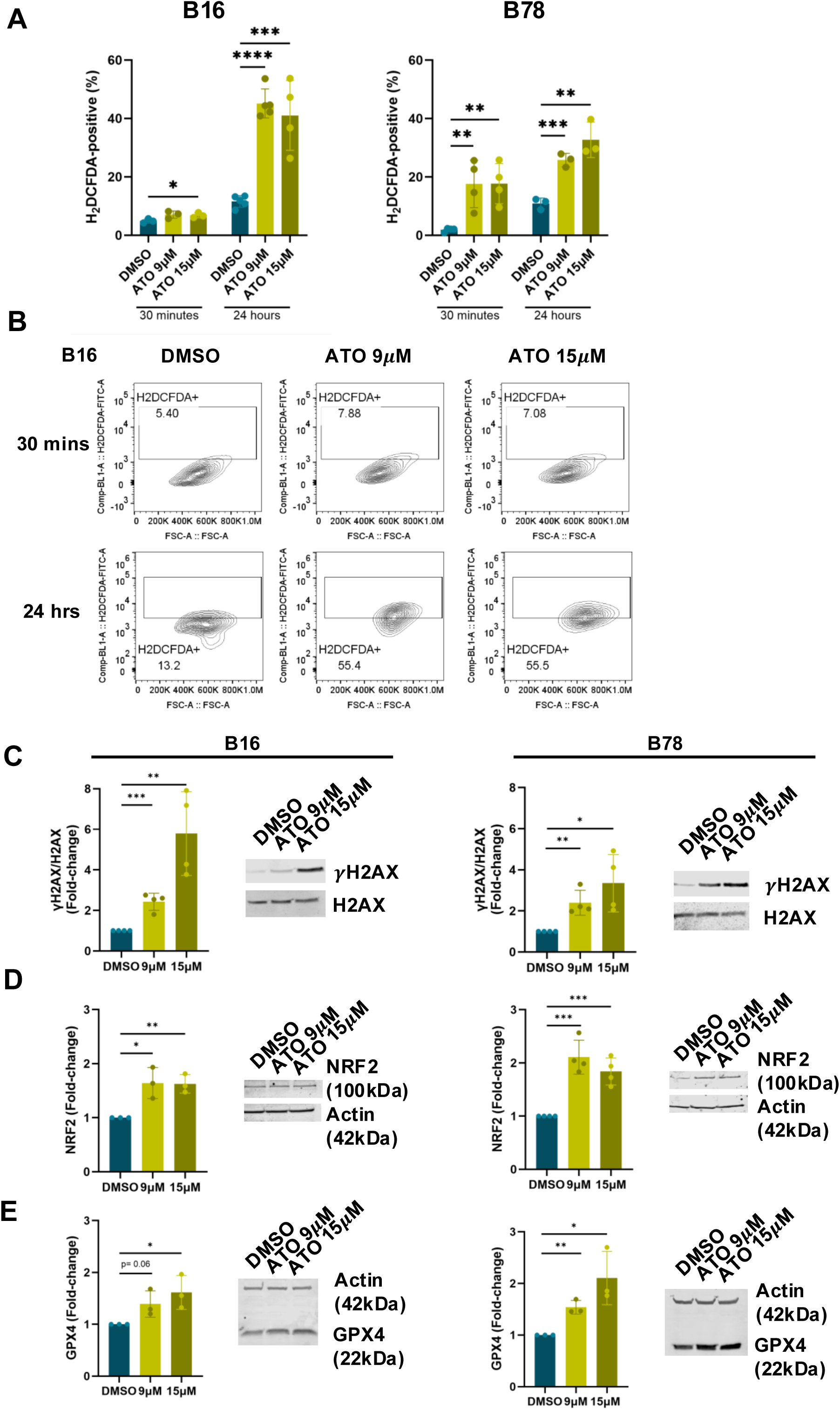
Atovaquone increases reactive oxygen species levels leading to oxidative stress in melanoma cells. Atovaquone-treated cells were stained with H_2_DCFDA, a ROS marker, which was then measured via flow cytometry. ROS levels were significantly increased at 30 minutes (A) and 24 hours (B) following atovaquone treatment. B16 representative flow cytometry plots are shown (B). At 24 hours post-atovaquone treatment, we observed increased γ-H2AX relative to H2AX via Western blots (C). Additionally, we found that antioxidant proteins NRF2 (D) and GPX4 (E) were upregulated at 24 hours post-atovaquone treatment via Western blots. \**P*<0.05, \*\**P*<0.01, \*\*\**P*<0.001, \*\*\*\**P*<0.0001.

Given these elevated ROS and multiphoton metabolic imaging analyses suggesting an oxidative environment, we hypothesized that atovaquone induced oxidative stress in melanoma cells. At 24 hours post-treatment, atovaquone-treated cells exhibited significantly increased phosphorylation of H2A histone family member X (γ-H2AX), a marker of double-stranded DNA breaks, compared to vehicle-treated cells, consistent with DNA damage (**Figure 3C**). Additionally, treated cells showed upregulation of nuclear factor erythroid 2-related factor 2 (NRF2) and glutathione peroxidase 4 (GPX4), key effectors of the oxidative stress response (**Figure 3D-E**). Altogether, these results strongly support sustained oxidative damage in melanoma cells following atovaquone treatment.

### Atovaquone exhibits antitumorigenic effects against melanoma

Given its disruption of cellular energetics and activation of oxidative stress, we hypothesized that atovaquone would exert cytotoxicity in melanoma cells. MTT assays demonstrated that atovaquone significantly decreased the viability of B16 and B78 melanoma cells in a dose-dependent manner with average IC_50_ concentrations of 15 μM at 24 hours (**Figure 4A-B**).

**Figure 4.**
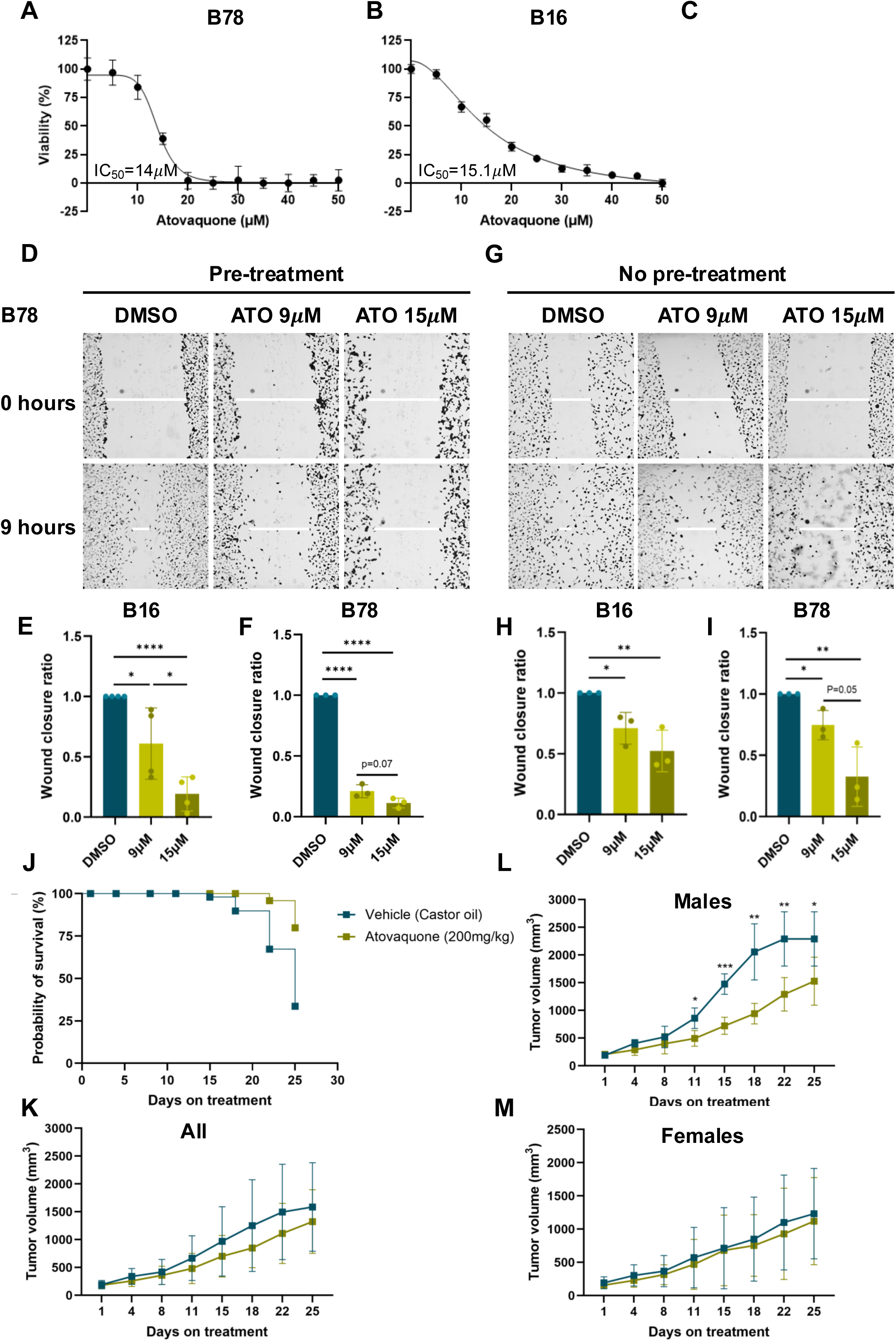
Atovaquone exhibits antitumorigenic effects in *in vitro* and *in vivo* melanoma models. MTT viability assays demonstrated dose-dependent effects of atovaquone in B16 and B78 cells at 24 hours post-treatment onset (A-B). Wound healing assays were performed using B16 and B78 cells. Cells were either pre-treated with atovaquone for 24 hours (A-B) or treated with atovaquone at wound induction (C-D). Wound width was calculated at 0 and 9 hours to calculate wound closure and compared to vehicle wound closure to obtain final ratio. Representative images of B78 cells are shown. Atovaquone-treated B78 mice showed increased survival rates in comparison to vehicle-treated mice (J). No differences in tumor volumes were observed in the overall cohort (K). However, atovaquone significantly decreased tumor volumes in atovaquone-treated male mice (N=4) when compared to vehicle-treated male mice (N=5) (L). No differences in tumor volumes were observed in female mice (Atovaquone N=8, Vehicle N=7) (M). \**P*<0.05, \*\**P*<0.01, \*\*\**P*<0.001, \*\*\*\**P*<0.0001.

To evaluate effects beyond cell viability, we assessed melanoma cell migration using wound healing assays. Atovaquone-treated cells exhibited significantly impaired wound closure compared to vehicle controls (**Figure 4D-F**). To account for viability-related confounding, we repeated the assay simultaneously introducing the wound and atovaquone treatment. Even under these conditions, atovaquone impaired wound closure (**Figure 4G-I**), indicating a direct effect on migration. These findings suggest that atovaquone impairs both cell viability and motility, highlighting its potential as a therapeutic agent for melanoma.

### Atovaquone controls melanoma tumor growth

Based on our *in vitro* findings, we next tested the *in vivo* efficacy of atovaquone using a B78 melanoma mouse model. Since atovaquone is administered orally, we performed pharmacokinetic (PK) studies to confirm its absorption into the bloodstream (**Supplemental Table 2).** Mice were treated for five days, and plasma was collected up to three days following treatment cessation. Average plasma concentrations were 93 μg/μl on day 1, 15 μg/μl on day 2, and 1 μg/μl on day 3. These concentrations were comparable to those seen in our atovaquone-treated ovarian cancer mouse models (data not shown) and consistent with steady-state concentrations in humans.^26^ These results confirm effective absorption and metabolic clearance of atovaquone in our B78 melanoma mouse model.

To evaluate the efficacy of atovaquone, mice received atovaquone suspension (200 mg/kg) or vehicle (castor oil) via oral gavage for four weeks. No treatment-related adverse effects were observed, consistent with atovaquone’s known low toxicity profile.^27^ At the end of the treatment period, atovaquone-treated mice exhibited significantly improved survival compared to vehicle controls (**Figure 4J**). These studies were conducted using both male and female mice randomized into treatment groups. Although no significant differences in overall tumor growth were initially observed, sex-specific analysis revealed that atovaquone-treated male mice had significantly reduced tumor volumes (**Figure 4K-M**).

### Atovaquone influences tumor metabolism

To investigate the metabolic effects of atovaquone in tumors, we performed optical metabolic imaging using multiphoton microscopy of *ex vivo* B78 tumors at early (post-treatment day 1) (PTD1) and late (post-treatment day 9) (PTD9) time points. Automated cell segmentation enabled single tumor cell quantification of NAD(P)H and FAD intensity and lifetime changes. This revealed striking metabolic heterogeneity with clusters of cells exhibiting different metabolic phenotypes within the same tumor (**Figure 5A**).

**Figure 5.**
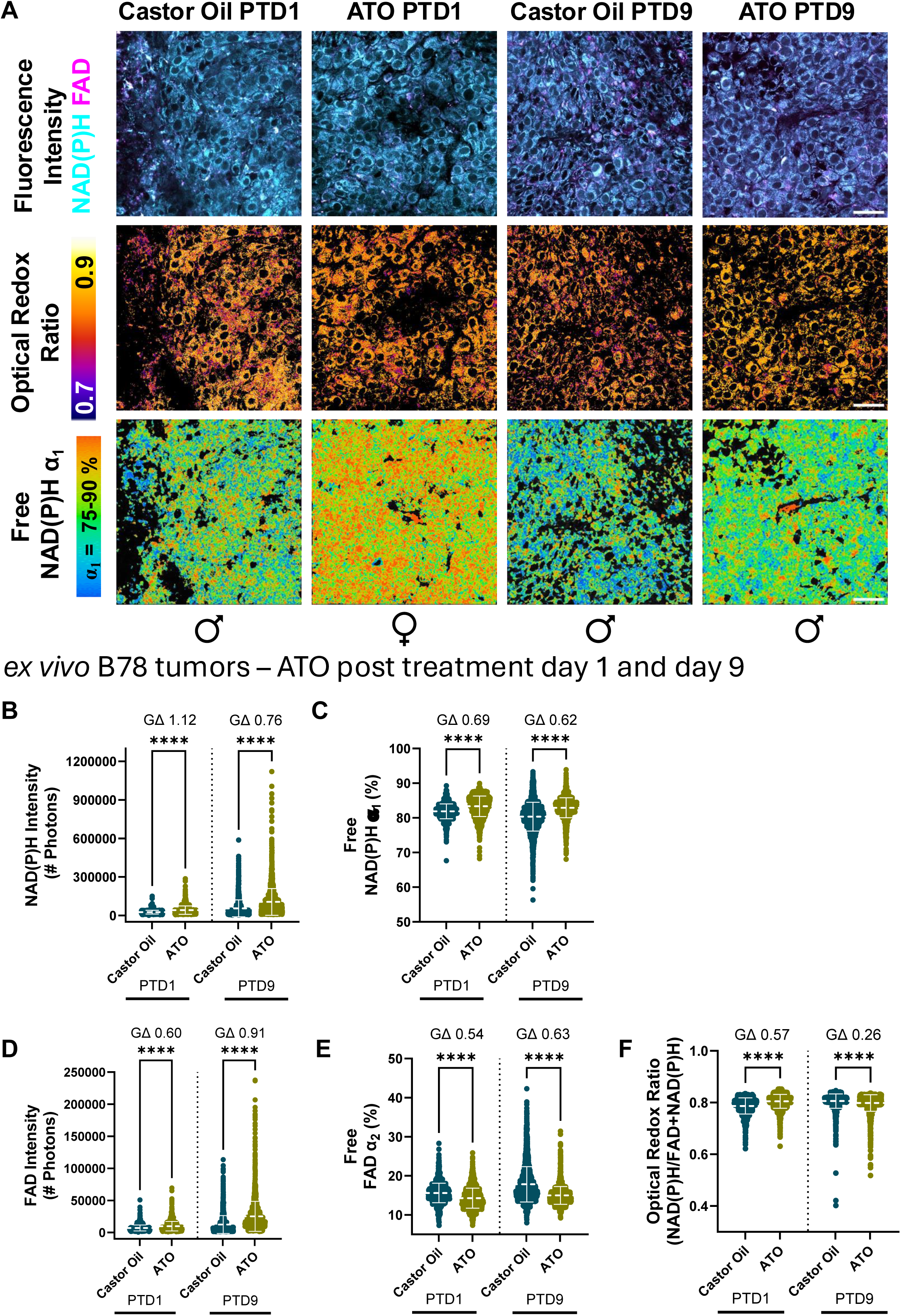
Optical metabolic imaging shows *ex vivo* B78 tumor metabolic changes during atovaquone treatment. (A) Representative *ex vivo* B78 tumor images showed fluorescence intensity changes of metabolic coenzymes NAD(P)H and FAD (top), optical redox ratio changes (middle), and NAD(P)H fluorescence lifetime changes (bottom) over time during atovaquone treatment (scale bar 50 μm). (B-F) Quantified single tumor cell B78 metabolic changes imaged at post-treatment day 1 (PT1D) and post-treatment day 9 (PTD9) (castor oil control, 200 mg / kg atovaquone in castor oil, *n* = 6 castor oil control mice, *n* = 6 atovaquone treated mice, *n* = 1175 castor oil control tumor cells PTD1, *n* = 1972 atovaquone treated tumor cells PTD1, *n* = 2513 castor oil control tumor cells PTD9, *n* = 2415 atovaquone treated tumor cells PTD9). \*\*\*\**P* < 0.0001, Glass’s Δ effect size, 2 biological replicates, mean ± SD.

At PTD1, both NAD(P)H and FAD fluorescence intensities were increased in atovaquone-treated tumors relative to vehicle controls **(Figure 5B,D**). Additionally, the ORR was elevated (**Figure 5F**), suggesting a proportionally greater increase in NAD(P)H relative to FAD, a reduced redox state, and a possible metabolic shift toward glycolysis. By PTD9, NAD(P)H and FAD intensities remained elevated, but the ORR was decreased in atovaquone-treated tumors (**Figure 5B,D,F**). These results indicate a greater relative increase in FAD, an oxidized redox state, and a possible shift towards more oxidative metabolism, similar to what was observed in our *in vitro* studies. Quantification of fluorescence lifetime changes revealed increased free NAD(P)H (α_1_) and decreased free FAD (α_2_) in atovaquone-treated tumors compared to vehicle controls at both time points (**Figure 5C, E**). These differences were more pronounced at day 9, underlying the ongoing evolving responses to metabolic pressures and oxidative stress. Overall, metabolic imaging highlighted complex metabolic changes occurring in *ex vivo* tumors following short (1 day) and long term (9 days) atovaquone treatment.

CD8-mCherry reporter mice were used in this study given our initial hypothesis stating that atovaquone might promote immune activation in melanoma. Metabolic imaging data suggested a trend toward increased number of intratumoral CD8+ T cells from atovaquone-treated mice compared to vehicle at PTD9 (**Figure S1A**). Additionally, we observed an increase in NAD(P)H intensity (significant at PTD1, trending at PTD9 *p* = 0.0844), free NAD(P)H α_1_ (significant at PTD1 and 9), and optical redox ratio (significant at PTD1 and 9) in intratumoral CD8+ T cells from atovaquone-treated mice at both timepoints (**Figure S1B-D**), suggesting a glycolytic phenotype, which is often associated with increased CD8+ T cell activation and function. Flow cytometry immunophenotyping also showed increased intratumoral CD8+ T cells in atovaquone-treated mice compared to vehicle at PTD10, and a trend towards an increase in CD8+ T cell cytokine expression, though no significant (**Figure S1E-G**).

### Atovaquone also exhibits effects in BRAF-mutated melanoma

To explore whether atovaquone’s antitumorigenic effects were observed in melanomas with different mutational profiles, we tested its effects on *BRAF*-mutated models, the most common somatic mutation observed in malignant melanoma.^28^ Using MTT viability assays, we observed that atovaquone exhibited cytotoxic effects in both human (MRA5, MRA6) and murine (YUMM1.7) *BRAF*-mutated melanoma cells (**Figure 6A-D**).^29,30^

**Figure 6.**
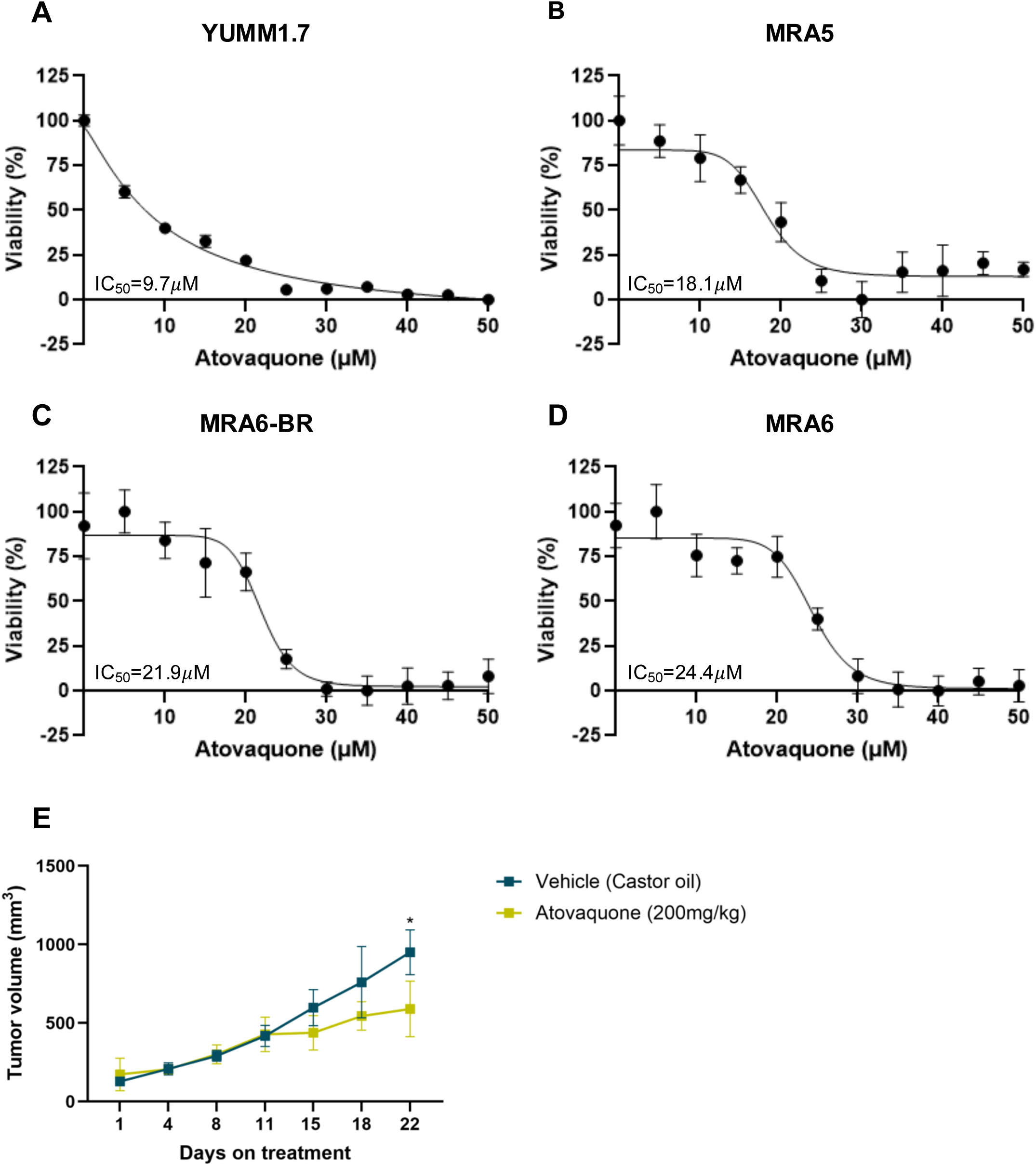
Atovaquone has antitumorigenic effects in BRAF-mutated melanoma models. MTT viability assays demonstrated dose-dependent effects of atovaquone in YUMM1.7, MRA5, MRA6, and MRA6-BR cells at 24 hours post-treatment (A-D). We observed a significant decrease in tumor volumes at day 22 post-treatment initiation in atovaquone-treated mice (*N*=4; 3 male, 1 female) compared to vehicle-treated mice (*N*=4; 2 male, 2 female). \**P*<0.05, \*\**P*<0.01, \*\*\**P*<0.001, \*\*\*\**P*<0.0001.

To evaluate its *in vivo* efficacy, we treated male and female mice bearing YUMM1.7 tumors with atovaquone suspension (200 mg/kg) or vehicle (castor oil). By day 22 post-treatment initiation, tumor growth was significantly reduced in the atovaquone-treated group compared to controls (**Figure 6E**). No significant survival benefits or sex differences were observed at these time points.

## DISCUSSION

Metabolic reprogramming is now recognized as a hallmark of cancer.^31,32^ Early studies by Warburg and colleagues suggested that tumors rely on aerobic glycolysis to support proliferation and growth.^33,34^ Since then, other metabolic pathways, including OXPHOS, have been identified as important contributors to tumor behavior. As tumors progress, metabolic phenotypes can diverge, influenced by many factors, such as nutrient and oxygen availability, cell-matrix and cell-cell interactions, and access to blood supply.^35–37^ Targeting these metabolic pathways has garnered increasing interest as a therapeutic strategy for cancer.

OXPHOS, responsible for approximately 70% of ATP in normal cells under physiologic conditions, remains active in many tumors, including melanomas.^38^ Although the rate of OXPHOS utilization varies by tumor type and stage of progression, most tumors, including melanomas, retain functional, and targetable, OXPHOS machinery.^7,38,39^ High OXPHOS expression and utilization have been associated with melanoma metastasis and chemoresistance.^40–42^ Our Seahorse data showed that OXPHOS contributes substantially to ATP production in melanoma cells accounting for 35-51% of the total ATP production rates. Additionally, multiphoton metabolic imaging revealed intratumoral metabolic heterogeneity that included cells with oxidative phenotypes. These findings provide support for targeting OXPHOS in melanoma.

Atovaquone, an FDA-approved antimalarial with a safe toxicity profile, has more recently attracted attention as a cancer therapeutic.^17–19,43,44^ In this study, we evaluated atovaquone in melanoma for the first time and demonstrated that it successfully inhibits OXPHOS, decreasing its contribution to ATP production to below 2%. While glycolytic rates were transiently increased following atovaquone treatment, this compensatory response was not sustained, leading to profound effects on total ATP levels. We have observed similar effects in epithelial ovarian cancer models treated with atovaquone, where both OXPHOS and glycolysis were downregulated at 24- and 48-hours post-treatment (unpublished data). Altogether, these results suggest that atovaquone targets OXPHOS in melanoma causing metabolic and energetic deficits that cells are unable to overcome.

Beyond its effects on metabolism, atovaquone also triggered oxidative stress in melanoma cells. Melanoma is considered a reactive oxygen species (ROS)-driven malignancy, characterized by elevated baseline ROS levels due to both intrinsic (e.g., melanin production) and extrinsic factors (e.g., ultraviolet radiation).^45^ In melanoma, enhancement of ROS-producing enzymes, growth factor signaling, and metabolic activity can further elevate ROS levels.^45–47^ ROS are usually maintained at subtoxic levels through robust antioxidant mechanisms. Strategies that aim to increase ROS levels to cytotoxic thresholds, overwhelming the antioxidant capacities and causing oxidative stress, may promote melanoma cell death.^47,48^ The ETC, involved in OXPHOS, is a major source of ROS, and its disruption leads to increased ROS.^24^ Thus, we suspected that atovaquone treatment would promote ROS accumulation. Flow cytometry analysis of the H_2_DCFDA ROS probe confirmed that ROS were elevated in atovaquone-treated cells compared to vehicle controls. Additionally, the changes in ORR obtained via metabolic imaging revealed that atovaquone-treated *in vitro* cells favored an oxidative state possibly due to increased ROS. Together with increased DNA damage and expression of antioxidant proteins, these results suggested that atovaquone treatment led to oxidative stress in melanoma cells.

These metabolic and oxidative effects ultimately led to antitumor outcomes. Atovaquone decreased cell viability with IC_50_ values in the low micromolar range, comparable to those observed in other solid tumors, such as epithelial ovarian cancer^19^ and breast cancer.^49^ Atovaquone also impaired wound closure, suggesting that mitochondrial metabolism could contribute to melanoma cell migration. This connection was also proposed in a different study showing that suppression of *PGC-1α*, a transcriptional coactivator of mitochondrial biogenesis, decreased cell dissemination and metastatic disease in B16 melanoma models.^50^ Our results contribute to growing evidence linking OXPHOS to both melanoma progression and invasion and highlight the impact of atovaquone on melanoma cell viability and migration.

Atovaquone significantly improved survival rates and impacted tumor growth in a B78 mouse model, particularly in male mice. While the sex-specific findings were unexpected, they may reflect previously reported differences in tumor growth rates between male and female C57BL/6 mice, including a study in a B16 melanoma model.^51–53^ In our B78 model, most vehicle-treated male mice reached the tumor volume humane end-point (2000 mm^3^) by day 25 (experimental end-point), while none of the female mice reached humane end-point volume by that time point. There are no reported sex differences in treatment outcomes or PK parameters (absorption and distribution) of atovaquone.^54^ Sex differences have been noted, however, in rodent studies using oil-based vehicles, attributed to differences in hormonal modulation of gut motility and absorption, body composition, and fat distribution.^55–57^ However, this is unlikely our case given that no sex differences were observed in the YUMM1.7 models, which were also developed in a C57BL/6 background and treated with the same oil-based ATO suspension. We will continue to consider and report on these sex differences in future studies.

Optical metabolic imaging of *ex vivo* B78 tumors further demonstrated the dynamic responses to atovaquone treatment. Both NAD(P)H and FAD intensities increased at early (PTD1) and late (PTD9) time points, indicating sustained metabolic disruption. However, the relative magnitude of these changes varied over time, resulting in an increased ORR at PTD1 and a decreased ORR at PTD9. Free NAD(P)H proportions were increased at both timepoints while free FAD proportions were decreased at both time points, with more pronounced changes on day 9. These shifts suggest an initial adaptation to metabolic disruption and oxidative stress that becomes dysregulated over time. These findings align with our *in vitro* data showing that atovaquone led to elevated ROS levels, decreased NAD(P)H intensity, and activation of antioxidant responses. These results support a model in which OXPHOS inhibition by atovaquone induces metabolic reprogramming and sustained oxidative stress in melanoma, ultimately saturating compensatory mechanisms and leading to cytotoxicity. Additionally, while flow cytometry and metabolic imaging data were suggestive of increased intratumoral CD8+ T cells in atovaquone-treated mice, further research is still needed to elucidate the immune effects of atovaquone in melanoma.

Ultimately, we examined whether atovaquone had activity across melanomas with different mutational profiles. Approximately 50-60% of melanomas have activating mutations in the *BRAF* gene, which have been linked to enhanced glycolysis.^28,58–60^ Despite these observations, we found that atovaquone impacted cell viability of *BRAF*-mutated lines and decreased tumor growth of a murine *BRAF*-mutated melanoma model. These findings suggest that atovaquone’s effects are consistent across different melanoma subtypes. Interestingly, BRAF inhibition has been shown to decrease glycolysis and increase mitochondrial mass and activity, and BRAF inhibitor-resistant melanoma models have been associated with elevated OXPHOS.^41,42^ Ongoing work is exploring the role of atovaquone in mitigating chemoresistance in melanoma.

## MATERIALS & METHODS

### 1. Cell lines and cell culture

Murine cell lines B16, B78, and YUMM1.7 were obtained from Drs. Paul Sondel (originally obtained from Ralph Reisfeld) and Vijay Setaluri at the University of Wisconsin, Madison. Human cell lines MRA2, MRA5, MRA6, and MRA6-BR were obtained from Dr. Mark Albertini at the University of Wisconsin, Madison.

Cells were maintained in the recommended culture media and incubated at 37°C in a humidified atmosphere with 5% CO_2_. Cell lines were tested for mycoplasma regularly using the Mycoscope^TM^ PCR Mycoplasma Detection Kit (Genlantis, Avantor, #10497-510). All experiments were performed on mycoplasma-negative cells.

### 2. Reagents and preparation

Atovaquone (purity ≥98%) was purchased from TCI America (Avantor, #A2545200MG) and solubilized in DMSO (ThermoFisher Scientific, #J66650.AP) at 10mg/ml. Stocks were kept at - 20°C and thawed as needed.

### 3. Cell viability assays

Cell viability was measured using 3-(4,5-dimethylthiazol-2-yl)-2,5-diphenyltetrazolium bromide (MTT) assays as previously described.^19,61^ Cells were seeded in 96-well plates and incubated overnight. They were then treated with stated agents or vehicle (DMSO). Following treatment, MTT (Sigma-Aldrich, #475989) solution (0.2mg/ml in PBS) was added to the wells, and cells were incubated at 37°C. After 3 hours, media containing MTT solution was removed and 100μl of DMSO was added to each well. Absorbance was measured at 570nm using the SpectraMax M3 Micro-Plate Reader at the Cancer Pharmacology Laboratory at the University of Wisconsin, Madison. ATP was measured using the CellTiter-Glo® 2.0 assay (Promega, #G9242). Cells were plated in opaque-walled 96-well plates and incubated overnight. They were then treated with atovaquone or vehicle for 24 hours. Equal volumes of CellTiter-Glo® 2.0 reagent were added to the wells and mixed with cell supernatants. Plates were incubated at room temperature for 10 minutes and luminescence was measured using the SpectraMax M3 Micro-Plate Reader at the Cancer Pharmacology Laboratory at the University of Wisconsin, Madison.

### 4. Western blots

Cells were washed with PBS and lysed in RIPA lysis buffer (ThermoFisher Scientific, #NC9839321) containing protease and phosphatase inhibitors. Protein concentrations were measured using the Pierce BCA Protein assay (ThermoFisher Scientific, #23227). Lysates equivalent to 25-30μg of total protein were electrophoresed, blotted to PVDF membranes, and probed with primary and secondary antibodies. Primary antibodies used were phosphorylated (γ-) H2AX (Cell Signaling Technology, #9718), H2AX (Cell Signaling Technology, #2595), NRF2 (Cell Signaling Technology, #12721), and GPX4 (Cell Signaling Technology, #59735). Beta actin (Santa Cruz Biotechnology, #sc-47778) was used as control. Secondary antibodies (IRDye), intercept antibody diluents, and blocking buffer were purchased from Licor (#926-32211, #926-68070, #927-65001, #927-60001). Protein bands were imaged and quantified using the Odyssey DLx imager.

### 5. Seahorse analysis

Oxygen consumption rates (OCR) and extracellular acidification rates (ECAR) were measured using a Seahorse XFe96 Analyzer. Cells were seeded in a 96-well plate supplied by the Seahorse XF Real-Time ATP Rate Assay Kit (Agilent, #103592-100) and incubated overnight at 37°C with 5% CO_2_. Cells were then treated with atovaquone (9 and 15µM) or vehicle (DMSO) for 1 or 24 hours for assessment at early and later time points. Cells were washed and incubated with atovaquone or vehicle in assay media for 1 hour at 37°C without CO_2_ to reach the ideal pH and temperature conditions as per the ATP Rate Assay protocol. The assay was run as detailed in the Agilent protocol. Oxygen consumption rate (OCR) and extracellular acidification rate (ECAR) readings were obtained. The readings were done after injections with glucose (10mM), followed by oligomycin (2µM), and then 2DG (100mM). After the assay, total protein content in each well was quantified using the BCA assay. ECAR and OCR values were normalized to the total protein content. Normalized ECAR and OCR values were used to calculate the percentage of the contribution of glycolysis and oxidative phosphorylation to ATP production using the Seahorse Wave Software (Agilent). These percentages were used to calculate the ATP rate index below.

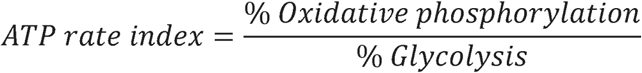

### 6. Migration assays

Cells were seeded in a 12-well plate and incubated overnight. Cells were pre-treated with atovaquone for 24 hours or treated at the time of wound. Wound was introduced (timepoint 0 hours) using a 20μl pipette tip. Cell migration was monitored via phase-contrast microscopy every hour until 9 hours post-scratch. Images were taken with a Leica Thunder microscope and analyzed using ImageJ software (RRID:SCR_003070). Width length of wound was calculated using the ‘Wound healing size tool updated’ Image J plug-in.^62^ Wound closure was calculated using the formula below and normalized to vehicle (DMSO) to obtain wound closure ratio.

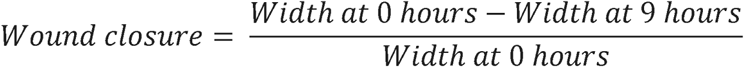

### 7. Reactive oxygen species assay

Following atovaquone treatment, cells were incubated with 10μM H_2_DCFDA (ThermoFisher Scientific, #D399) in PBS for 45 minutes. Cells were washed and resuspended in buffer (2% FBS in PBS) containing DAPI (2μg/ml). Fluorescence minus one (FMO), unstained, and positive (H_2_O_2_) controls were used. Data was obtained using the ThermoFisher Attune flow cytometer at the Flow Cytometry Laboratory at the University of Wisconsin Carbone Cancer Center and analyzed using FlowJo software.

### 8. *In vivo* studies

#### 8.1. B78 mouse model

Institutional Animal Care and Use Committee (IACUC) approval (M006730) was obtained prior to conducting animal experiments. Mice were housed at the Small Animal Facility at the University of Wisconsin Carbone Cancer Center.

Adult male and female C57BL/6 mice were injected intradermally in the right flank with 2×10^6^ B78 or YUMM1.7 cells resuspended in 100 µl of PBS. Once tumors reached 100 mm^3^, we began treatment with 100µl of either atovaquone suspension (200 mg/kg) or vehicle control (castor oil) delivered via oral gavage five days per week (Monday-Friday) for four weeks. Atovaquone concentration was selected according to previous reports on the use of atovaquone for the treatment of solid tumors.^19,63^ Tumor sizes (width and length) were measured regularly with calipers to calculate tumor volumes using the formula below. Mice were euthanized at the experiment endpoint (four weeks on treatment), or if they reached humane endpoint (tumors > 2000mm^3^, significant weight loss, or other signs of distress). We compared tumor volumes and survival rates between the control and treatment groups.

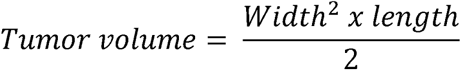

For metabolic imaging studies, CD8-mCherry reporter mice were obtained from Dr. Melissa Skala at the Morgridge Institute for Research. The CD8α-mCherry knock-in mouse was designed and generated by the University of Wisconsin Genome Editing and Animal Model cores. The mCherry-P2A knock-in cassette resulted in a CD8 reporter allele, rather than an mCherry-CD8 fusion. A sequence-confirmed founder was identified, the founder was backcrossed to C57BL/6J to produce F1 progeny, and F1 progeny were similarly sequence-confirmed. Full details have been previously described.^21,64^

#### 8.2. Pharmacokinetic (PK) studies

PK studies were performed after one week (5 days) of atovaquone treatment. Plasma was collected at days 1, 2, and 3 post-treatment cessation. Plasma from untreated mice was used as negative control and paclitaxel was used as the internal standard. Separations were obtained on Shimadzu HPLC System with Waters Symmetry C18 column (150mm length x 3.9mm inner diameter, 5 μm particles). The injections carried 5 μl of samples. The mobile phase was composed of a mixture of acetonitrile (85% v/v) and 0.02M potassium phosphate buffer (pH 5.5, 15%v/v). The flow rate was set at 1.0 mL/min. The detection was carried on Shimadzu Diode Array Detector SPD-M20A at 254nm. HPLC assay and analysis were performed by the Cancer Pharmacology Laboratory at the University of Wisconsin Carbone Cancer Center.

#### 8.3. Flow cytometry immunophenotyping

Tumors were extracted and processed into single cell suspensions following published protocols.^65^ Cells were first stained with Zombie NIR^TM^ Fixable Dye (Biolegend, #423106) for 15 minutes at room temperature. Without washing, surface antibodies were added and incubated for 30 minutes on ice. Cells were then fixed (4% paraformaldehyde) for 20 minutes. Intracellular antibodies were added in permeabilization buffer (0.1% saponin) and incubated for 30 minutes on ice. Antibody titrations were performed prior to experiments. Specific antibody clones and volumes used are detailed in **Supplemental Table 3**. Data was collected using the ThermoFisher Attune flow cytometer at the Flow Cytometry Laboratory at the University of Wisconsin Carbone Cancer Center and analyzed using FlowJo software.

### 9. Multiphoton Imaging

#### 9.1. Ex vivo B78 tumors from CD8-mCherry mice

*Ex vivo* imaging of B78 tumors in CD8-mCherry mice was performed in vehicle castor oil treated mice (*n* = 6) and 200 mg / kg atovaquone treated mice (*n* = 6) across days 1 and 9 post-treatment. Tumors were well established (∼200-800 mm^3^), 5-7 weeks after inoculation. Imaging was performed as two separate biological replicates. *Ex vivo* imaging of whole B78 tumors was performed immediately following euthanasia via CO_2_. Excised tumors were secured to an imaging dish with PBS coupling and tape. *Ex vivo* imaging was complete within one to two hours post mouse euthanasia, accurately capturing cellular metabolism based on our previous work that showed that *ex vivo* metabolism is statistically identical to *in vivo* metabolism for up to 12 hours post euthanasia.^66^

Autofluorescence images were captured with a custom-built multi-photon microscope (Bruker) using an ultrafast femtosecond laser (Ultrafast Coherent Chameleon Ultra II, InSight DS+). Fluorescence lifetime measurements were performed using time-correlated single photon counting electronics (Becker & Hickl). Fluorescence emission, in FLIM mode, was detected simultaneously in three channels using bandpass filters of 466/40 nm (NAD(P)H), 514/30 nm (FAD), and 650/45 nm (mCherry) prior to detection with three GaAsP photomultiplier tubes (Hamamatsu). All three fluorophores were simultaneously excited using a previously reported wavelength mixing approach.^21,67,68^ During wavelength mixing in FLIM mode, typical power at the power meter was 1.4-2.5 mW for the 750 nm laser and 1.5-3.0 mW for the 1041 laser. All images were acquired with a 40×/1.13 NA water-immersion objective (Nikon) at 512×512pixel resolution and an optical zoom of 1.0-2.0. NAD(P)H and FAD intensity and lifetime images were acquired to sample metabolic behavior of B78 tumor and CD8 T cells across 8-16 fields of view and multiple depths within each tumor.

#### 9.2. In vitro B78 tumor cells

*In vitro* imaging of B78 tumor cells was performed by plating 100,000-150,000 tumor cells in RPMI-1640 media, on coated glass imaging dishes (2 technical replicates) 48 hours prior to imaging. To inhibit OXPHOS / electron transport chain complex 3, cells were first imaged untreated, then treated with 15 μM atovaquone (Ato) in castor oil and DMSO, then imaged again at either 1 hour or 24 hours later. To inhibit fatty acid oxidation, cells were first imaged untreated, then treated with 100 nM etomoxir (Eto) in PBS, then imaged again 24 hours later. To inhibit pentose phosphate pathway, cells were first imaged untreated, then treated with 5 mM 6-aminonicotinamide (6AN) in DMSO, then imaged again 30 minutes later. B78 cells were imaged twice across two biological replicates for each inhibitor treatment.

Autofluorescence images were captured with a custom-built multi-photon microscope (Bruker) using an ultrafast femtosecond laser (Ultrafast Coherent Chameleon Ultra II, InSight DS+). The laser was tuned to 750 nm for NAD(P)H excitation and tuned to 890 nm for FAD excitation. The average power at the power meter was ∼2-4.5 mW for NAD(P)H excitation and ∼4-4.7 mW for FAD excitation. A pixel dwell time of 4.8 μs was used for all images. NAD(P)H and FAD images were acquired sequentially. A 440/80 nm bandpass filter isolated NAD(P)H emission onto the photomultiplier tube (PMT) detector. A dichroic mirror directed wavelengths greater than 500 nm onto a 550/100 nm bandpass filter, isolating FAD emission onto a second PMT. Fluorescence lifetime measurements were acquired with time-correlated single photon counting electronics (Becker and Hickl) and a GaAsP PMT (Hamamatsu). All images were acquired with a 40×/1.13 NA water-immersion objective (Nikon) at 512×512 pixel resolution and an optical zoom of 1.0. NAD(P)H and FAD intensity and lifetime images were acquired to sample metabolic behavior of B78 tumor cells across 4-5 fields of view per dish, per condition.

#### 9.3. Multiphoton Imaging Analysis

##### Fluorescence Lifetime Fitting

The fluorescence lifetimes of free and protein-bound NAD(P)H and FAD are distinct, and these lifetimes along with their weights can be recovered with a two-exponential fit function. Therefore, fluorescence lifetime decays for both NAD(P)H and FAD were fit to the following bi-exponential function in SPCImage:

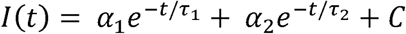

For NAD(P)H, τ_1_corresponds to the free lifetime, τ_2_ corresponds to the protein-bound lifetime, and the weights (α_1_, α_2_; α_1_ + α_2_ = 1) correspond to the proportion of free and protein-bound NAD(P)H, respectively.^69–72^ Conversely for FAD, τ_1_ corresponds to the protein-bound lifetime and τ_2_corresponds to the free lifetime.^22,71,73^ An instrument response function was measured using SHG (900 nm excitation) from urea crystals for input into the decay fit procedure. The following fluorescence lifetime endpoints were calculated from the fitted model: τ_1_, τ_2_, α_1_, and α_2_ for both NAD(P)H and FAD; along with the optical redox ratio.^70,72,74^

##### Cell Segmentation

Automated cell segmentation was performed through Cellpose, with manual corrections made as needed using the Napari viewer through a custom Python script.^75,76^ Single tumor cells were identified, circled, and segmented as whole cells (nuclei and cytoplasm captured) using NAD(P)H intensity images. The resulting segmented images were saved as masks. Using these masks and the raw imaging data, fluorescence lifetime variables and redox ratio values were calculated for each individual cell using custom code adapted from Cell Analysis Tools.^77^ Fluorescence lifetime variables were calculated at a pixel level for the *in vitro* data, and on a region of interest or cell level, with a single decay curve for each individual cell, for the *ex vivo* data.^78^ Calculations were performed using Python / Spyder.

### 10. Statistical analysis

Statistical analysis for the methodology described above was done using GraphPad Prism (V10.0.2) software. A minimum sample size of 3 biological replicates was used to determine significance. Significance thresholds were placed at *P*-values <0.05.

For *ex vivo* tumor data, ordinary one-way ANOVA with Sidak’s multiple comparisons test was used to assess differences in OIM parameters between treatment groups. For *in vitro* cell data, Mann–Whitney statistical tests for non-parametric, unpaired comparisons were performed to assess differences in OMI parameters between treatment groups. Effect size was also calculated to assess differences in optical metabolic imaging parameters between treatment groups using Glass’s Δ because comparisons of very large sample sizes of individual cells nearly always pass traditional significance tests unless the population effect size is truly zero. Glass’s Δ is defined as: (mean experimental group – mean control group) ⁄ standard deviation control group. A Glass’s Δ > 0.8 was chosen to indicate significant effect size based on previous studies.^21,79,80^ Results are represented as dot plots showing mean ± standard deviation where each dot represents a single tumor cell (GraphPad Prism 10.4).

## Ethics Statement

This study was performed in accordance with the Declaration of Helsinki. Collection of animal tissue samples for this study was approved as part of the study protocol. This animal study was approved by Institutional Animal Care and Use Committee at the University of Wisconsin School of Medicine and Public Health.

## Data Availability Statement

The data generated in this study are available within the article and its supplementary data files.

## Acknowledgments

We would like to thank Alexander Fedorov and the University of Wisconsin Carbone Cancer Center (UWCCC) Cancer Pharmacology Laboratory for their contributions, which included the PK studies, as well as the University of Wisconsin School of Medicine and Public Health Department of Obstetrics and Gynecology and the Wisconsin Ovarian Cancer Alliance for the support provided. We also acknowledge the UWCCC Flow Cytometry Laboratory and the Biomedical Research Model Services for the use of their facilities and services. The following funding sources are also acknowledged: The Carol Skornicka Chair in Biomedical Imaging; Retina Research Foundation Daniel M. Albert Chair; the National Institutes of Health [R01CA278051, R01CA272855, R01CA238423-05, R01CA238423-04S1]; and the United States Department of Veterans Affairs [IO1BX005627].

## Authors Contribution Statement

- Mayra Betancourt Ponce: conceptualization, data curation, formal analysis, investigation, methodology, resources, validation, visualization, writing – original draft preparation
- Alexa R. Heaton: conceptualization, data curation, formal analysis, investigation, methodology, resources, software, validation, visualization, writing – review and editing
- Samantha Burkard: investigation, formal analysis
- Aurora D’Amato: investigation, formal analysis
- Nicha Boonpattrawong: investigation, formal analysis
- Melissa Skala: funding acquisition, project administration, supervision, software, resources, writing – review and editing
- Manish Patankar: conceptualization, funding acquisition, methodology, project administration, resources, supervision, writing – review and editing
- Lisa Barroilhet: conceptualization, funding acquisition, methodology, project administration, resources, supervision, writing – review and editing

## Supplementary Materials

**Supplemental Table 1. Seahorse analysis of melanoma cells following atovaquone treatment.**

**Supplemental Table 2. Pharmacokinetic studies of B78 mice treated with atovaquone.**

**Supplemental Table 3. Flow cytometry immunophenotyping panel**

**Supplemental Table 4. Optical metabolic imaging parameters and their descriptions.**

**Supplemental Figure 1. CD8+ T cells tend to be increased in atovaquone-treated B78 mice. (**A) Representative *ex vivo* B78 tumor images showed fluorescence intensity changes of metabolic coenzymes NAD(P)H (blue) and FAD (green) plus mCherry+ CD8 T cells (red) infiltrating the tumors over time during atovaquone treatment (scale bar 50 μm). (B-D) Quantified single CD8 T cell metabolic changes imaged at post-treatment day 1 (PT1D) and post-treatment day 9 (PTD9) (castor oil control, 200 mg / kg atovaquone in castor oil, *n* = 6 castor oil control mice, *n* = 6 atovaquone treated mice, *n* = 383 castor oil control CD8 T cells PTD1, *n* = 211 atovaquone treated CD8 T cells PTD1, *n* = 44 castor oil control CD8 T cells PTD9, *n* = 150 atovaquone treated CD8 T cells PTD9). Glass’s Δ effect size, 2 biological replicates, mean ± SD. (E) A significant increase in intratumoral CD8+ T cells (CD45+CD3+CD8+) was observed in atovaquone-treated mice compared to vehicle at PTD10. Percentages of CD8+ T cells in treatment groups were normalized to vehicle controls. (F-G) No significant changes were observed in the percentage of CD8+ T cells expressing cytokines IFN-γ and TNF-α. Flow cytometry *N*=4 mice per treatment group. \**P*<0.05, \*\**P*<0.01, \*\*\**P*<0.001, \*\*\*\**P*<0.0001.

