## Supplemental Tables 1-4, Supplemental Figure 1 for "Atovaquone-induced therapeutic rewiring of melanoma metabolism"

**Supplemental Table 1**. Seahorse analysis of melanoma cells following atovaquone treatment.

| Groups | | GlycoATP Production Rate (pmol/min) | | MitoATP Production Rate (pmol/min) | | Total ATP Production Rate (pmol/min) | | XF ATP Rate Index | | % Glycolysis | | % OXPHOS | |
|---|---|---|---|---|---|---|---|---|---|---|---|---|---|
| | | Avg | SD | Avg | SD | Avg | SD | Avg | SD | Avg | SD | Avg | SD |
| B16 | DMSO (1h) | 771.8 | 174.5 | 642.2 | 204.2 | 1414.0 | 374.3 | 0.82 | 0.10 | 55.1 | 3.2 | 44.9 | 3.2 |
| | ATO (9uM; 1h) | 1104.4 | 329.2 | 17.0 | 9.9 | 1121.4 | 335.4 | 0.016 | 0.007 | 98.5 | 0.7 | 1.5 | 0.7 |
| | ATO (15uM; 1h) | 956.5 | 325.8 | 9.0 | 5.2 | 965.6 | 330.6 | 0.009 | 0.003 | 99.1 | 0.3 | 0.87 | 0.28 |
| | DMSO (24h) | 669.7 | 96.6 | 415.3 | 101.8 | 1085.0 | 194.6 | 0.61 | 0.08 | 62.1 | 3.1 | 37.9 | 3.1 |
| | ATO (9uM; 24h) | 285.5 | 69.9 | 2.8 | 3.4 | 288.3 | 70.3 | 0.010 | 0.012 | 99.0 | 1.1 | 0.98 | 1.13 |
| | ATO (15uM; 24h) | 290.2 | 98.9 | 1.7 | 2.0 | 291.9 | 98.9 | 0.006 | 0.007 | 99.4 | 0.7 | 0.61 | 0.69 |
| B78 | DMSO (1h) | 347.4 | 73.9 | 364.9 | 90.1 | 712.3 | 157.5 | 1.0 | 0.1 | 49.1 | 3.6 | 50.9 | 3.6 |
| | ATO (9uM; 1h) | 630.5 | 107.2 | 6.9 | 2.5 | 637.5 | 107.3 | 0.011 | 0.005 | 98.9 | 0.5 | 1.1 | 0.5 |
| | ATO (15uM; 1h) | 744.0 | 72.9 | 6.1 | 3.1 | 750.0 | 74.0 | 0.008 | 0.004 | 99.2 | 0.4 | 0.81 | 0.40 |
| | DMSO (24h) | 336.7 | 94.0 | 283.6 | 229.4 | 620.3 | 298.3 | 0.76 | 0.60 | 64.1 | 25.2 | 35.9 | 25.2 |
| | ATO (9uM; 24h) | 290.4 | 47.4 | 2.6 | 3.3 | 293.0 | 49.3 | 0.008 | 0.011 | 99.2 | 1.0 | 0.81 | 1.04 |
| | ATO (15uM; 24h) | 131.0 | 51.5 | 2.2 | 3.2 | 133.2 | 50.1 | 0.024 | 0.041 | 97.8 | 3.7 | 2.2 | 3.7 |

**Supplemental Table 2.** Pharmacokinetic studies of B78 mice treated with atovaquone.

| Samples | IS Retention Time | IS Area | ATO Retention Time | ATO Area | Area ATO/IS | Calculated µg/mL | Average (µg/mL) |
|---|---|---|---|---|---|---|---|
| Day 1 | 1.26 | 156,000 | 3.75 | 266,200 | 1.71 | 91.96 | **93** |
| Day 1 | 1.26 | 154,200 | 3.75 | 267,500 | 1.73 | 93.47 | |
| Day 2 | 1.26 | 152,200 | 3.80 | 39,800 | 0.26 | 14.83 | **15** |
| Day 2 | 1.26 | 152,500 | 3.80 | 40,700 | 0.27 | 15.12 | |
| Day 3 | 1.26 | 156,700 | 3.88 | 1,000 | 0.01 | 1.21 | **1** |
| Day 3 | 1.26 | 154,700 | 3.88 | 1,100 | 0.01 | 1.25 | |
| Control | 1.26 | 158,800 | 3.80 | 0 | 0.00 | 0.00 | |
| Control | 1.26 | 591,000 | 3.80 | 0 | 0.00 | 0.00 | 0 |
| Control | 1.26 | 504,900 | 3.80 | 0 | 0.00 | 0.00 | |
| Control | 1.26 | 506,500 | 3.80 | 0 | 0.00 | 0.00 | 0 |

IS= Internal Standard (Paclitaxel)
All samples run in duplicates
Calculation parameters: Slope= 0.0187, Y-interval= -0.0163

**Supplemental Table 3.** Flow cytometry immunophenotyping panel

| Marker | Fluorophore | Clone | Volume (µl per $10^6$ cells) |
|---|---|---|---|
| CD45 | FITC | 30-F11 | 1 |
| CD3 | SparkBlue574 | 17A2 | 2 |
| CD4 | BV510 | GK1.5 | 5 |
| CD8 | Alexa594 | 53-6.7 | 1 |
| CD11c | BV711 | N418 | 0.15 |
| CD25 | PE-Cy7 | PC61 | 2.5 |
| NKp46 | BV605 | 29A1.4 | 2.5 |
| FoxP3 | Alexa647 | MF-14 | 2 |
| TNFα | PE | MP6-XT22 | 0.625 |
| IFNγ | BV421 | XMG1.2 | 2.5 |
| Live/Dead | NIR | | 0.5 |

| **Supplemental Table 4.** Optical metabolic imaging parameters and their descriptions. | |
|---|---|
| Imaging Parameter | Description |
| NAD(P)H $\tau_1$ | Free NAD(P)H lifetime |
| NAD(P)H $\tau_2$ | Protein-bound NAD(P)H lifetime |
| NAD(P)H $\alpha_1$ | Proportion of free NAD(P)H |
| NAD(P)H $\alpha_2$ | Proportion of protein-bound NAD(P)H |
| NAD(P)H $\tau_m$ | NAD(P)H mean lifetime: $\alpha_1 \tau_1 + \alpha_2 \tau_2$ |
| FAD $\tau_1$ | Protein-bound FAD lifetime |
| FAD $\tau_2$ | Free FAD lifetime |
| FAD $\alpha_1$ | Proportion of protein-bound FAD |
| FAD $\alpha_2$ | Proportion of free FAD |
| FAD $\tau_m$ | FAD mean lifetime: $\alpha_1 \tau_1 + \alpha_2 \tau_2$ |
| Optical redox ratio | ORR: Intensity NAD(P)H / (Intensity NAD(P)H + Intensity FAD) |

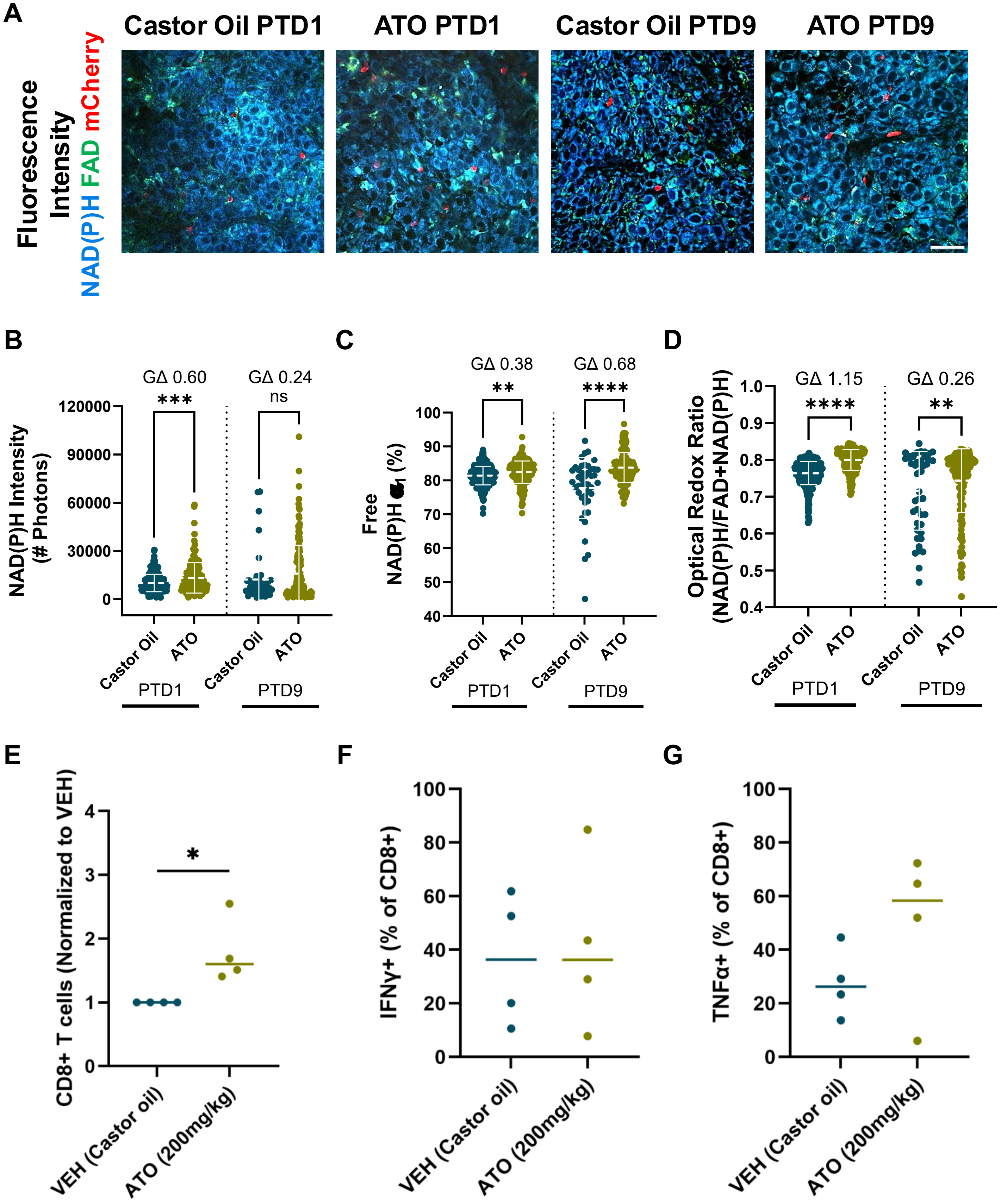
A
Castor Oil PTD1
ATO PTD1
Castor Oil PTD9
ATO PTD9
Fluorescence Intensity
NAD(P)H FAD mCherry
B
GΔ 0.60
***
GΔ 0.24
ns
NAD(P)H Intensity (# Photons)
120000
90000
60000
30000
0
Castor Oil
ATO
PTD1
PTD9
C
GΔ 0.38
**
GΔ 0.68
****
Free NAD(P)H α1 (%)
100
90
80
70
60
50
40
D
GΔ 1.15
****
GΔ 0.26
**
Optical Redox Ratio (NAD(P)H/FAD+NAD(P)H)
1.0
0.9
0.8
0.7
0.6
0.5
0.4
E
*
CD8+ T cells (Normalized to VEH)
4
3
2
1
0
VEH (Castor oil)
ATO (200mg/kg)
F
IFNγ+ (% of CD8+)
100
80
60
40
20
0
G
TNFα+ (% of CD8+)
